# Nrf2 attenuates the innate immune response after experimental myocardial infarction

**DOI:** 10.1101/2022.01.10.475615

**Authors:** Daniel I. Bromage, Silvia Trevelin, Josef Huntington, Victoria X. Yang, Ananya Muthukumar, Sarah J. Mackie, Greta Sawyer, Xiaohong Zhang, Celio X.C. Santos, Niloufar Safinia, Ioannis Smyrnias, Mauro Giacca, Alex Ivetic, Ajay M. Shah

## Abstract

**Objectives:** We aimed to investigate the contribution of the transcription factor nuclear factor erythroid-derived 2-like 2 (Nrf2) to the inflammatory response after experimental myocardial infarction (MI).

**Background:** There is compelling evidence implicating dysregulated inflammation in the mechanism of ventricular remodeling and heart failure (HF) after MI. The transcription factor Nrf2 (encoded by *Nfe2l2*) is a promising target in this context. It impedes transcriptional upregulation of pro-inflammatory cytokines and is anti-inflammatory in various murine models.

**Methods:** We subjected Nrf2^-/-^ mice and wild type (WT) controls to permanent left coronary artery (LCA) ligation. The inflammatory response was investigated with fluorescence-activated cell sorting (FACS) analysis of peripheral blood and heart cell suspensions, together with qRT-PCR of infarcted tissue for chemokines and their receptors. To investigate whether Nrf2-mediated transcription is a dedicated function of leukocytes, we interrogated publicly available RNA-sequencing (RNA-seq) data from mouse hearts after permanent LCA ligation for Nrf2-regulated gene (NRG) expression.

**Results:** FACS analysis demonstrated a profoundly inflamed phenotype in the hearts of global Nrf2^-/-^ mice as compared to WT mice after MI. Moreover, infarcted tissue from Nrf2^-/-^ mice displayed higher expression of inflammatory cytokines, chemokines, and their receptors, including IL6, *Ccl2*, and *Cxcr4*. RNA-seq analysis showed upregulated NRG expression in WT mice after MI compared to untreated mice, which was significantly higher in bioinformatically isolated CCR2^+^ cells.

**Conclusions:** Taken together, the results suggest that Nrf2 signalling in leukocytes, and possibly CCR2^+^ monocyte-derived cardiac resident macrophages, may be potential targets to prevent post-MI ventricular remodeling.

## Introduction

Despite timely reperfusion, myocardial infarction (MI) is the commonest cause of heart failure (HF), and mortality remains high in such patients (1–5). HF after MI is caused by ventricular remodeling, characterised by fibrosis, ventricular dilatation, and progressive functional deterioration. Post-MI inflammation is vital to remove non-viable cells and activate reparative mechanisms to prevent myocardial rupture. Myeloid cells are especially important in regulating inflammation after MI. The balance between inflammatory and reparative subsets appears crucial and there is compelling evidence implicating excessive inflammation in the mechanism of remodeling. (6, 7).

In the traditional paradigm, ischaemic tissue-resident macrophages contribute to neutrophil and inflammatory monocyte recruitment (8). Next, inflammatory Ly6C^hi^ monocyte-derived M1 macrophages secrete pro-inflammatory cytokines, including IL-6, that mediate apoptosis and matrix degradation (6, 9–13). These macrophages later shift to a reparative M2 phenotype, secrete anti-inflammatory cytokines, and instigate myocardial repair (14–16). Single cell (sc) RNA-seq experiments have implicated cardiac resident macrophages (RMs) in mediating the inflammatory response to cardiac injury (17–19). C-C chemokine receptor 2-expressing (CCR2^+^) RMs (derived from recruited monocytes) and CCR2^-^ RMs (maintained independent of monocyte recruitment) promote and inhibit, respectively, monocyte recruitment after MI (20). However, leukocyte signalling after MI is not fully understood.

One promising candidate pathway involves the transcription factor nuclear factor erythroid-derived 2-like 2 (Nrf2). Nrf2 regulates a network of cytoprotective genes (Nrf2-regulated genes, NRGs) and appears to reduce infarct size in models of ischaemia-reperfusion injury (21–24). Additionally, global Nrf2 induction and deletion protect against and exacerbate, respectively, remodeling after MI with permanent left coronary artery (LCA) ligation (25, 26). Previous studies have linked the protective effects of Nrf2 to its antioxidant actions (27); however, Nrf2 has been shown to have anti-inflammatory effects in other settings. Nrf2 also impedes transcriptional upregulation of pro-inflammatory cytokines in M1 macrophages (28), and both global and leukocyte-specific Nrf2 deletion exacerbate inflammation in various murine models, including sepsis and emphysema (29–33). The gene encoding Nrf2 (*Nfe2l2*) has been shown by scRNA-seq to be enriched in inflammatory macrophages of murine atherosclerotic aortas, but there was also clear expression in other subsets, indicating a complex contribution to leukocyte biology (34). The role of Nrf2 in regulating the inflammatory response after MI is unknown. The aim of this study was to investigate the contribution of Nrf2 in leukocytes to the inflammatory response after MI.

## Methods

Detailed methods are provided in Supplemental methods. All use of animals was in accordance with the United Kingdom (Scientific Procedures) Act 1986 (PPL70/8889) and institutional guidelines. Permanent LCA or sham ligation was performed on adult female Nrf2^-/-^ mice and Nrf2^fl/fl^ (WT) littermates both on a C57BL/6J background (35). Nrf2 expression was assessed by Western blot, immunofluorescence, and qRT-PCR of canonical Nrf2-regulated genes. Infarct size was quantified as a proportion of area at risk after 24Ͱh of recovery. In separate experiments, qRT-PCR was used to quantify mRNA expression in the infarct region after 72 h of recovery, and fluorescence-activated cell sorting (FACS) of blood and heart digests was performed to examine the cellular inflammatory phenotype.

For RNA sequencing analysis, NRGs were defined according to published chromatin immunoprecipitation sequencing (ChIP-seq) experiments using models of constitutive nuclear accumulation (Keap1^-/-^) or depletion (Nrf2^-/-^) of Nrf2 (36). To look for an NRG signature in leukocytes, these were cross-referenced with publicly available bulk RNA-seq data from various adult cardiac cell populations from mice subjected to permanent LCA ligation or sham surgery (GEO accession GSE95755 (37)). Gene ontology (GO) analysis was performed with MetaCore™. To investigate whether the Nrf2-regulated response is a function of specific leukocyte subsets, we used publicly available scRNA-seq data of CD45^+^ cells taken from infarcted tissue 4d after MI, chosen to align with FACS experiments (GEO accession GSE106473 (38)). Unbiased clusters were generated and identified using the Immunological Genome (ImmGen) compendium and canonical subset markers. We calculated an NRG score from the summed expression of the 10 most highly expressed NRGs.

Results were compared using a two-tailed t-test or Mann-Whitney U test (non-parametric) for 2 groups of continuous variables, and analysis of variance (ANOVA) and Tukey’s Multiple Comparison Test for 3 or more groups. Data is presented as meanͰ±ͰSEM, and p<0.05 was considered significant.

## Results

### Global Nrf2^-/-^ mice display excessive leukocyte recruitment after MI

Global Nrf2^-/-^ mice have been phenotypically well characterized under baseline conditions (39, 40). To assess the effects of Nrf2 independent of confounding changes in infarct size, we employed a permanent ligation model. After 24 h of experimental LCA ligation, Nrf2 protein levels were significantly increased compared to sham ligation in WT mice **(Supplemental Fig. 1A)**. Heart macrophages (CD68^+^ cells) in the infarcted area showed higher nuclear translocation of Nrf2 after MI, which was lost in Nrf2^-/-^ mice **(Supplemental Fig. 1B)**. Accordingly, hearts of Nrf2^-/-^ mice displayed diminished expression of *Nfe2l2* and canonical Nrf2-regulated genes in the heart **(Supplemental Fig. 1C-D)**. Infarct size as a proportion of area at risk was no different between WT and Nrf2^-/-^ mice at 24 h (WT 93.0±1.3% vs. Nrf2^-/-^ 88.3±7.9%, **Supplemental Fig. 1E-G)**.

Next, we investigated the inflammatory response to MI. We chose a time point of 72 h, which has been shown to coincide with peak monocyte infiltration (41). The infarct zone from WT and Nrf2^-/-^ mice **(Fig 1A)**, or a respective area of left ventricle (LV) from sham controls, and peripheral blood samples were analysed by FACS (gating strategy, **Fig. 1B-C)**. While no differences were apparent in blood at this timepoint **(Supplemental Fig. 2)**, there was a profoundly enhanced inflammatory response in infarcted tissue of Nrf2^-/-^ compared to WT mice **(Fig. 1)**. This comprised total leukocytes (defined as CD45^+^) and key myeloid subsets, including neutrophils (CD45^+^CD19^-^Ly6G^+^) and monocytes (CD45^+^CD19^-^Ly6G^-^Ly6C^hi^) **(Fig. 1D-E)**. Furthermore, there was a more pro-inflammatory (CD206^-^, N1) than proresolution (CD206^+^, N2) neutrophil phenotype. However, there was no significant expansion of monocyte-derived or -independent RMs (defined as CD45^+^Ly6C^lo^CCR2^+^ and CCR2^-^, respectively). Intracellular staining demonstrated increased protein expression of inflammatory cytokines and chemokines, IL-6, iNOS, CD11b and CCR2, in cardiac macrophages **(Fig. 1F)**. This is consistent with the finding that infarcted Nrf2^-/-^ hearts also had increased mRNA expression of the chemokine (C-C motif) ligand 2 (*Ccl2/Mcp1*) and the chemokine receptor, *Cxcr4* **(Fig. 1F)**, which contributes to leukocyte recruitment to the infarcted region. Taken together, these results indicate an inhibitory function of Nrf2 on the inflammatory response after experimental MI.

**Fig. 1:**
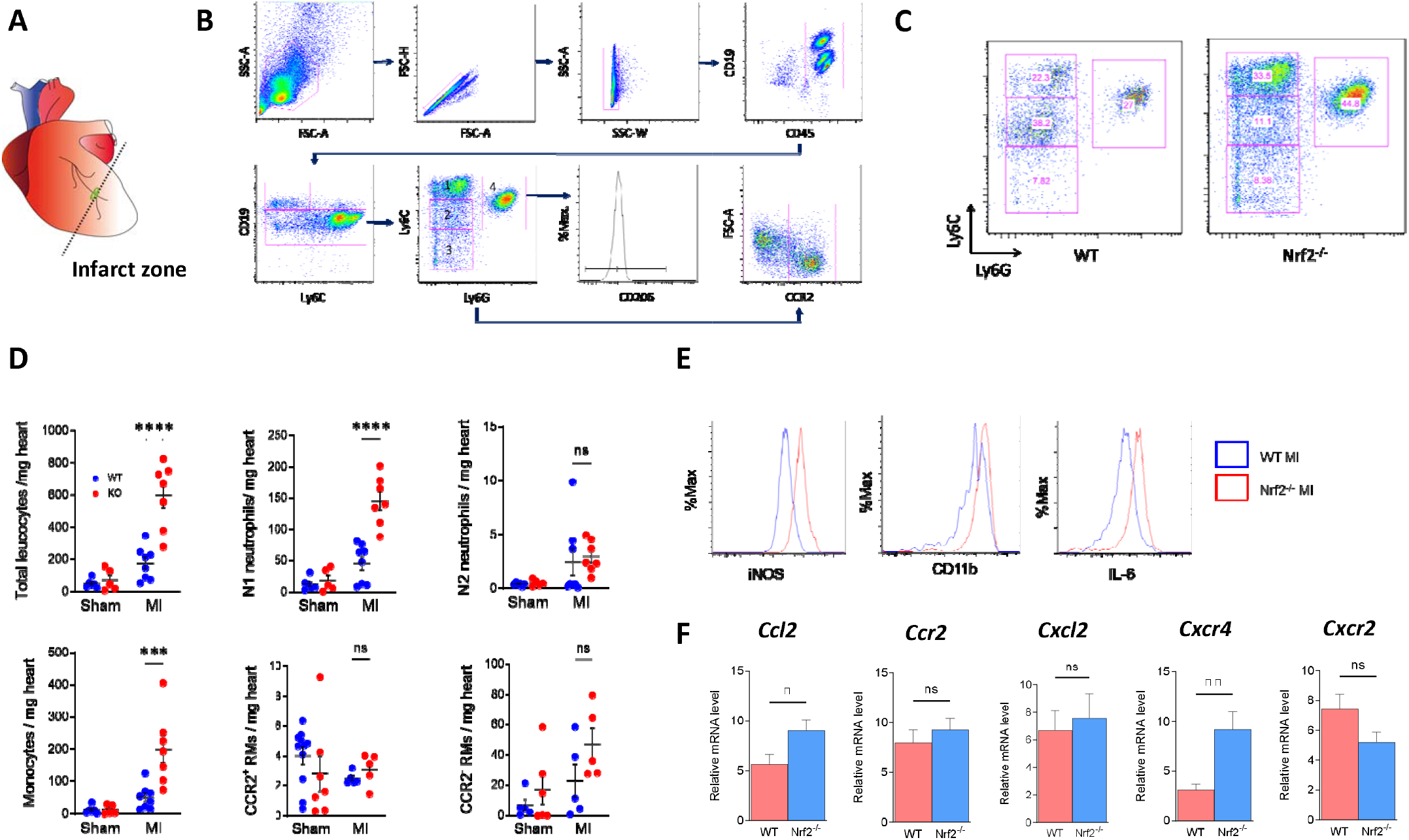
Nrf2^-/-^ mice have a pro-inflammatory phenotype after MI. 9-13 week old mice were anaesthetised and subjected to permanent LAD ligation prior to recovery for 72 h. **A.** Schematic of MI experiment in which the infarct zone was isolated from excised hearts; **B.** Gating strategy for FACS analysis (1-3, Ly6C^hi^, Ly6C^lo^ and Ly6C^-^, respectively, monocytes and macrophages; 4, neutrophils); **C.** Representative plots from FACS analysis showing higher leukocyte infiltration in Nrf2^-/-^ mice after MI; **D**. Quantification of leukocytes in heart digests, defined as (left to right): CD45^+^(total leukocytes), CD45^+^CD19^-^ Ly6G^+^CD206^-^ (N1) and CD45^+^CD19^-^Ly6G^-^CD206^-^ (N2) neutrophils (top), and monocytes (CD45^+^CD19^-^Ly6G^-^Ly6C^hi^), CCR2^+^ RMs (CD45^+^CD19^-^Ly6G^-^Ly6C^lo^CCR2^+^), and CCR2^-^ RMs (CD45^+^CD19^-^Ly6G^-^Ly6C^lo^CCR2^-^) (bottom). Data shown as cells/mg, n=5-8 per group compared using 1-way ANOVA with Tukey’s post-test; **E.** Representative histograms from Ly6C^hi^ cells showing expression of inflammatory modulators, iNOS, CD11b, and IL-6; **F.** Expression in the infarct region in WT mice (n=10), compared to Nrf2^-/-^ mice (n=12-13). Data presented as mean ± SEM and compared using a t-test. *p<0.05, **p<0.01, ***p<0.001.

### Differential NRG expression in leukocytes after experimental MI

We next wished to determine whether NRGs are relevant in leukocytes themselves. For this, publicly available bulk RNA-seq data from adult mice on day 4 after permanent LCA ligation or sham surgery was analysed (38). Selected results from this dataset were first validated by performing qRT-PCR on our own experimental infarct tissue obtained at day 3 after permanent LCA occlusion or sham in WT mice, finding increased expression of *Hmox1, Il10, Il6, Sod2* and *Prdx3* compared to sham **(Fig. 2A)**. We next cross-referenced gene expression data with published ChIP-seq data for Nrf2 targets (36) and found 209 differentially expressed NRGs in leukocytes, compared to 45, 60, and 24 in cardiac myocytes, fibroblasts, and endothelial cells, respectively **(Fig. 2B)**. Clustering of pathway enrichment analysis using gene ontology (GO) annotations highlighted canonical pathways in leukocytes relating to regulation of fibrosis via the TGF-β superfamily **(Fig. 2C)**. This is consistent with the concept that leukocyte NRG expression contributes to the regulation of ventricular remodeling after MI.

**Fig. 2:**
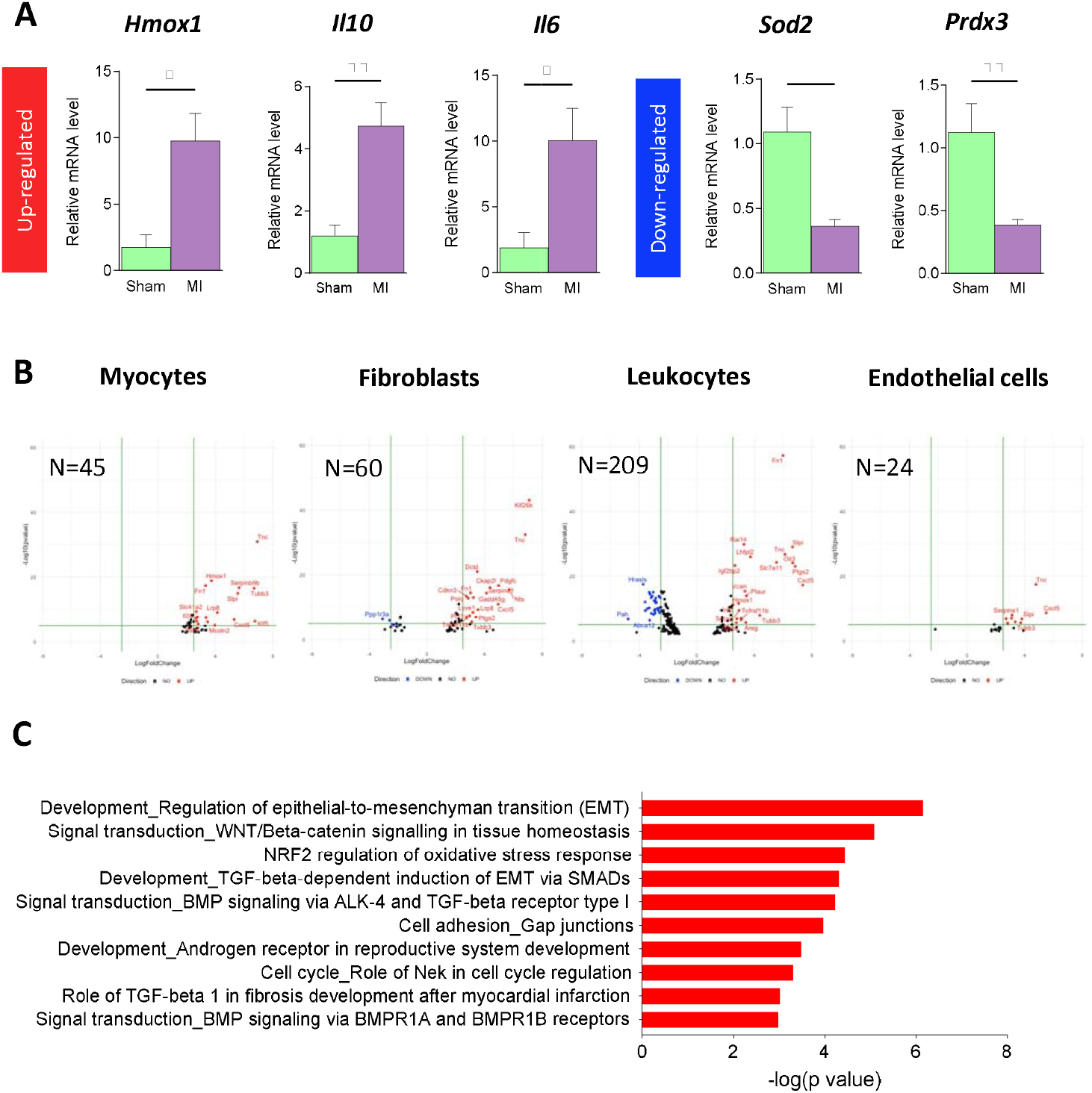
Myocardial infarction activates Nrf2-dependent signalling in leukocytes. **A**. Validation of publicly available bulk RNAseq data; *Hmox1, Il10, Ccl2, Cxcr2, Ccr2, Sod2, Prdx3* and *Il6* expression in the infarct region on D3 after MI (n=9-10), compared to sham in WT mice (n=4-6). Data presented as mean ± SEM and compared using a t-test. *p<0.05, **p<0.01, ***p<0.001; **B.** Volcano plots showing differentially expressed Nrf2-regulated genes in different cell types day 3 after MI or sham surgery in adult mice (using publicly available data). N represents the number of differentially expressed Nrf2-regulated genes; **C**. Gene enrichment analysis of GO pathways relating to Nrf2-regulated genes.

### NRG expression is a feature of CCR2^+^ cardiac monocytes and macrophages

We used publicly available scRNA-seq data from n=1,858 CD45^+^ sorted leukocytes taken from the infarcts of 4 WT mice at day 4 after MI, and n=703 unsorted cells from the corresponding area of a single, non-infarcted WT heart (38). Unbiased clustering of MI leukocytes generated 11 clusters, which were identified using ImmGen and canonical markers **(Fig. 3A; Supplemental Figs. 3-14)** and annotated for *Nfe2l2* expression **(Fig. 3B)**. Based on the observation that cardiac RMs are central role to leukocyte recruitment after MI, an NRG score was calculated from the summed expression of the most highly expressed NRGs in bioinformatically isolated cardiac monocytes and macrophages (*Fth1, Ctsb, Ftl1, Dusp1, Mpeg1, Lcp1, Ubc, Hmox1, Fn1, Cebpb*). The global NRG score, applied to MI and WT datasets, was significantly increased in MI compared to untreated mice (p<0.0001, **Fig. 3C)**. Further analysis of NRG expression revealed significantly higher expression in CCR2^+^ monocytes and macrophages (p<0.0001, **Fig. 3D)**. Expression of individual NRGs according to cell cluster are shown in **Supplemental Fig. 15**.

**Fig 3:**
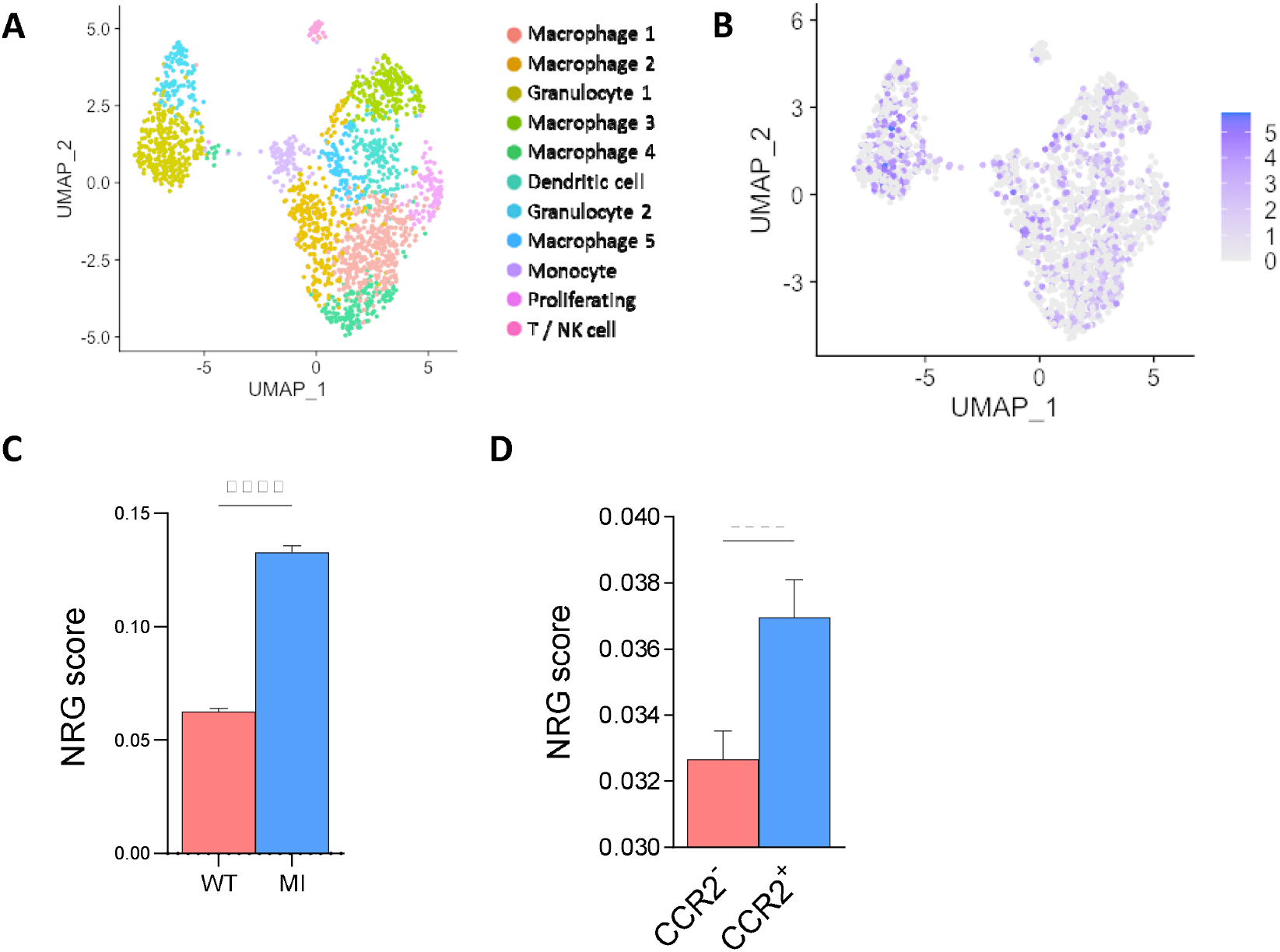
Nrf2-regulated gene (NRG) expression in CCR2+ cardiac monocytes and macrophages. **A**. Publicly available single-cell RNA-seq data for 1,858 single cells isolated from a single WT mouse 4d after permanent LCA occlusion, clustered and displayed on a uniform manifold approximation and projection (UMAP) plot. Cell lineage inferred by marker gene signatures; **B**. UMAPs annotated with Nfe2l2 expression; **C**. Global Nrf2-regulated gene (NRG) score based on summed expression of NRGs in MI and WT datasets, normalized to cell number; **D**. NRG score in CCR2^-^ and CCR2^+^ monocytes and macrophages, normalized to cell number. Data presented as mean ± SEM and compared using the Mann-Whitney test, ****p<0.0001.

## Discussion

In summary, we demonstrate a robust pro-inflammatory phenotype in global Nrf2^-/-^ mice after MI, encompassing significantly increased pro-inflammatory leukocytes, cytokines, leukocyte-recruitment chemokines, and their receptors. Analysis of bulk RNA-seq data confirmed differential NRG expression in leukocytes after MI in WT mice, suggesting that Nrf2 contributes to the inflammatory response, and indicated functions relating to fibrosis and myocardial remodelling. More detailed analyses of scRNA-seq data showed upregulation of NRGs in MI compared to WT samples, which was especially pronounced in CCR2^+^ monocytes and macrophages.

The prominent inflammatory phenotype in Nrf2^-/-^ mice after MI was not confined to any leukocyte subset and comprised both neutrophils and macrophages. Previous studies have associated MI in global Nrf2^-/-^ mice with worse ventricular remodeling and LV function (26); however, this has previously been related to antioxidant and cytoprotective effects (27), and the extent and cellular composition of the inflammatory phenotype has not been described before, despite the known roles of Nrf2 in mediating inflammation (28, 42). Furthermore, hearts of Nrf2^-/-^ mice had higher expression of chemokines and chemokine receptors. This included higher *Cxcr4* but not *Cxcr2* mRNA expression, which may suggest aged neutrophils with a highly reactive phenotype (43), or recruitment of other CXCR4-expressing leukocyte subsets via stromal derived factor-1α (SDF-1α) expression (44, 45).

Using RNA-seq data, we confirmed *Nfe2l2* and NRG expression in leukocytes, and identified more differentially expressed NRGs compared to cardiac myocytes, fibroblasts, and endothelial cells. Our finding of upregulated RGs after MI is consistent with previous studies (25). We also observed significantly higher NRG expression in CCR2^+^ monocytes and macrophages, which may indicate an important role in monocyte-derived RMs. This contrasts with previous studies that suggest NRG expression is limited to tissue-resident CCR2^-^ cardiac RMs after MI (42). This may relate to how NRGs are selected and indicate distinct functions in diverse subsets. This is relevant because CCR2^+^ and CCR2^-^ cells have respectively negative and positive effects in several models of myocardial injury, including MI, and understanding the contribution of Nrf2 signalling may reveal novel therapeutic targets.

These data support the concept that Nrf2 suppresses inflammation, possibly by impeding over-recruitment of inflammatory cells. Targeting Nrf2 is challenging because it is expressed in several cell types and studies suggest the timing of Nrf2 upregulation is crucial, with most indicating a protective effect of acute expression but harmful effects at later timepoints (46, 47). Therefore, deciphering signaling pathways in the heart, together with identifying novel NRG-expressing leukocyte subsets, may identify novel therapeutic targets.

Persistently high mortality among patients with post-MI HF necessitates novel approaches. Several clinical trials evaluating non-specific immunosuppression have failed to ameliorate HF after MI (48), which is unsurprising given the complex relationship between inflammatory and reparative processes. Approaches specifically targeting detrimental inflammation are required. The CANTOS trial showed that targeting atherosclerosis with canakinumab, an anti-IL-1β monoclonal antibody, reduced major adverse cardiac events in high-risk patients (49). After MI, the anti-inflammatory agent colchicine reduced ischaemic cardiovascular events compared to placebo (50). These provide the first evidence of targeted approaches improving outcomes in cardiovascular disease. Our findings are clinically relevant as we demonstrate that Nrf2 in leukocytes has a fundamental role in cardiac inflammation after MI. Therefore, targeting acute inflammation via Nrf2-regulated pathways may be one such approach to preventing HF after MI.

## Acknowledgements

The authors wish to thank Matteo Beretta for his assistance with confocal imaging.

