## Supplemental appendix for "Nrf2 attenuates the innate immune response after experimental myocardial infarction"

### Supplemental methods

#### Genetically modified mice and experimental MI

All use of animals was in accordance with the United Kingdom (Scientific Procedures) Act 1986 (PPL70/8889) and institutional guidelines. Mice with global deletion of Nrf2 (Nrf2^-/-^) on a C57BL/6J background have been described previously (1, 2). Permanent LCA or sham ligation was performed on female Nrf2^-/-^ and wild type (WT) mice aged 9-13 weeks (3). Briefly, mice were anaesthetised with 2% isoflurane/98% oxygen and ventilated via orotracheal intubation. A lateral thoracotomy was made in the fourth intercostal space. The pericardium was removed and the LCA ligated 1–2 mm below the tip of the left atrium. The chest wall was repaired in layers. Mice recovered in a warmed chamber for at least 6 h. Intramuscular buprenorphine and subcutaneous flunixin was administered for perioperative analgesia.

#### Evaluation of infarct size

After 24 h of recovery, mice were sacrificed, and the heart removed for evaluation of infarct size. Hearts were cannulated and manually perfused with ice-cold phosphate buffered saline (PBS) until the effluent ran clear. The area at risk (AAR) was defined by perfusion of 200 μl Evans blue dye. Samples were frozen for 20 min at −80 °C and stained with triphenyltetrazolium chloride (TTC) for assessment of infarct size by slicing the heart into five 1 mm sections and incubating them for 20 min in the dark in 1% TTC in phosphate buffer. Following incubation, the sections were fixed in 10% formalin for 24 h before being scanned for analysis by planimetry using ImageJ (version 1.45s, NIH). Infarct size is expressed as a percentage of the AAR (IS/AAR).

#### Western blot

After 24 h of recovery, mice were sacrificed, and the heart excised. Heart tissues were homogenized in lysis buffer (composition: 10 mM HEPES pH 7.9, 50 mM NaCl, 0.5 M sucrose, 0.1 mM EDTA, 0.5% Triton X-100) containing phosphatase inhibitors (10 mM tetrasodium pyrophosphate, 100 mM NaF, 17 mM βglycerophosphate, 1 mM sodium orthovanadate), a protease inhibitor cocktail (Sigma) and a proteasome inhibitor (2 µg/ml Mg132). Protein content was estimated by a Bradford Assay. Equal amounts of lysates or protein extracts were loaded onto SD polyacrylamide gels, and then transferred to nitrocellulose membranes. After blocking the membranes with Tris-buffered saline and 0.05% Tween 20 (TBST) containing 5% non-fat milk for 1 hr at RT, the membranes were incubated overnight with primary anti-NRF2 antibody (Santa Cruz) at 4 ^o^C. After incubating with horseradish peroxidase (HRP)-linked secondary antibodies, the blots were revealed by chemiluminescence. β-actin (Sigma) was used as loading control. Densitometric analysis was performed using Image J software (NIH, USA). All values are presented as mean AU±SEM.

#### qRT-PCR

Hearts were arrested in diastole with KCL 72 h after MI or sham surgery, extracted and washed in ice-cold PBS. The infarcted region was visually dissected and snap frozen. mRNA was extracted using a dedicated kit (RNeasy, Qiagen) according to the manufacturer's instructions. Purified mRNA was converted to cDNA using the Omniscript RT Kit (Qiagen), as per the manufacturer's instructions. The qRT-PCR reaction was performed with SYBR Green and the comparative Ct method, using three reference genes (GAPDH, β-actin and CANX) (4). All primers were purchased from Sigma (see **Supplemental Table 1** for primer sequences).

#### Immunofluorescence

Sections from WT and Nrf2^-/-^ MI experiments described above were mounted in OCT before being cut into 5 μm sections at −20°C in a microtome-cryostat, transferred to slides, and dried at RT for 2 h. Sections were fixed with ice-cold methanol for 2 min, before blocking with 5% goat serum/PBS for 1 h at RT. Immunofluorescent co-staining of macrophages and Nrf2 was performed using 1:500 rat monoclonal anti-CD68 (MCA1957) and 1:100 rabbit polyclonal anti-Nrf2 (ab137550) diluted in 5% goat serum/PBS, overnight at 4°C. Anti-rabbit Alexa Fluor 488 and anti-rat Alexa Fluor 647 secondary antibodies (Invitrogen) and incubated at 1:400 dilution in the same buffer for 60 min at room temperature. DAPI nuclear stain (1:200) was added with the secondary antibodies to all sections. Coverslips were mounted using fluorescence mounting medium (Dako). After drying, Alexa 488 and Alexa 647 fluorescence was imaged using a 40× oil immersion objective, by sequential scanning using a Leica SP5 confocal microscope.

#### Fluorescence-activated cell sorting

In a separate experiment, blood was taken by direct cardiac puncture using a 23G needle and hearts were extracted after 72 h and washed with ice-cold PBS. Single-cell suspensions were prepared from heart tissue by digestion in a mixture of collagenase (1 mg/ml, Sigma-Aldrich), DNase (160 IU/ml), and hyaluronidase (500 IU/ml) at 37°C for 30 minutes. Samples were triturated and filtered through a 40-μm nylon mesh. Red blood cells were lysed in 2% NH_4_Cl buffer. Non-specific interactions were blocked with anti-mouse Fc block (1:50, BioLegend) before staining for 30 min at 4⁰C. For intracellular staining, the Foxp3 Transcription Factor Staining Buffer Set (eBioscience, 00-5523-00) was used. Full minus one (FMO) samples were prepared for IL6, CCR2, iNOS and CD206, in addition to an unstained control. Antibodies for surface and intracellular staining are described in **Supplemental Table 2**. Samples were acquired in an LSRFortessa flow cytometer (BD Biosciences) and analyzed using FlowJo software 9.7.5. Macrophages were identified as either CD45^+^CD19^-^Ly6G^-^Ly6C^hi^ (M1) or CD45^+^CD19^-^Ly6G^-^Ly6C^lo^ (M2), neutrophils were identified as either CD45^+^CD19^-^Ly6G^+^CD206^-^ (N1) or CD45^+^CD19^-^Ly6G^+^CD206^+^ (N2).

#### RNA sequencing analysis

NRGs were defined according to published chromatin immunoprecipitation sequencing (ChIP-seq) experiments using models of constitutive nuclear accumulation (Keap1^−/−^) or depletion (Nrf2^−/−^) of Nrf2 (5). These were cross-referenced with publicly available bulk RNA-seq data from adult (postnatal day 56; P56) mice day 4 after permanent LCA ligation or sham surgery (6). This data contains significantly differentially expressed genes (false discovery rate ≤0.05, log2(fold change) ≥1 or ≤-1), which were used to identify candidate NRGs (GEO accession GSE95755). Differential gene expression testing was performed in the original publication using EdgeR (6). Gene ontology and pathway analysis was applied to NRGs using MetaCore^TM^, and processes and maps, respectively, ranked according to their -log(p value).

To investigate whether the Nrf2-regulated response is a function of specific leukocyte subsets, we analysed publicly available scRNA-seq data (GEO accession GSE106473; (7)) using the Seurat 2.3.4 package in R as suggested by the accompanying tutorial (satijalab.org). The deposited data was already bioinformatically cleaned to remove ribosomal, haemoglobin and mitochondrial RNAs. Genes were removed if found in less than 3 cells, and empty droplets or doublets were removed by filtering barcodes that registered less than 200, or more than 2500 genes, respectively. Remaining cells were scaled to a total 1 × 10^4^ total molecules per cell and the data was log normalised.

The gene space was limited to over-dispersed high-variability genes. Linear dimension reduction was performed with principal-component analysis. Only principal components with significant variability were used for clustering, identified using the *JackStraw* function. Unsupervised clustering was performed on the significant PCs according to the default Seurat implementation of a community detection approach. Marker genes and associated *P* values were determined by differential expression using the Seurat implementation of the likelihood-ratio test based on zero-inflated data. Cell and cluster proximity were visualized by projecting onto a 2D space using Uniform Manifold Approximation and Projection (UMAP). Clusters were visualized using heat maps of clusters on x axis and marker genes on y axis, and violin plots depicting expression probability densities for each cluster.

To generate an NRG score from the single cell datasets, we cross-referenced the highest expressed NRGs in bioinformatically isolated monocytes and macrophages (clusters 0, 1, 3, 4, 7 and 8) with published NRGs to produce a final list of 10 (5). To define single-cell NRG scores, we summed counts for the constituent genes per cell and normalised to cell number by dividing individual cell NRG scores by the number of cells within its respective cluster.

#### Statistical analyses

Results were compared using a two-tailed t-test or Mann-Whitney U test (non-parametric) for 2 groups of continuous variables, and analysis of variance (ANOVA) and Tukey's Multiple Comparison Test for 3 or more groups . The Shapiro-Wilk test was performed to determine if data was normally distributed. Data is presented as mean ± SEM. Statistical significance was reported if P < 0.05 using the following nomenclature: *p<0.05, **p<0.01, ***p<0.001 and ****p<0.0001. Analyses were performed with GraphPad Prism® version 8.4.3 for Windows.

### Supplemental figures

#### Supplemental Fig 1: Characterisation of WT and Nrf2^-/-^ mice after MI

Mice were anaesthetised and subjected to permanent LAD ligation prior to recovery for 24h. **A.** The infarct area from WT mice was analysed by Western blot compared to sham operated mice with respect to Nrf2 protein expression; **B.** Immunofluorescent staining of the infarct region demonstrating nuclear translocation (arrowed) of Nrf2 (green) in relation to CD68 macrophages (red). The nucleus is stained with DAPI (blue). Representative images from n=3 independent experiments; Expression of *Nfe2l2* (**C**) and canonical Nrf2-regulated genes (**D**) after MI in WT (n=10) or Nrf2^-/-^ (n=13) mice subjected to MI; **E.** Analysis of their respective areas at risk, to ensure surgical consistency; **F.** IS as a proportion of AAR analysed using Evans Blue and TTC staining. **G.** Representative scanned transverse heart sections demonstrating Evans Blue area (blue), area at risk (pink) and infarct (white). Statistical significance was assessed using unpaired t tests, n=3-7. Data presented as mean ± SEM.

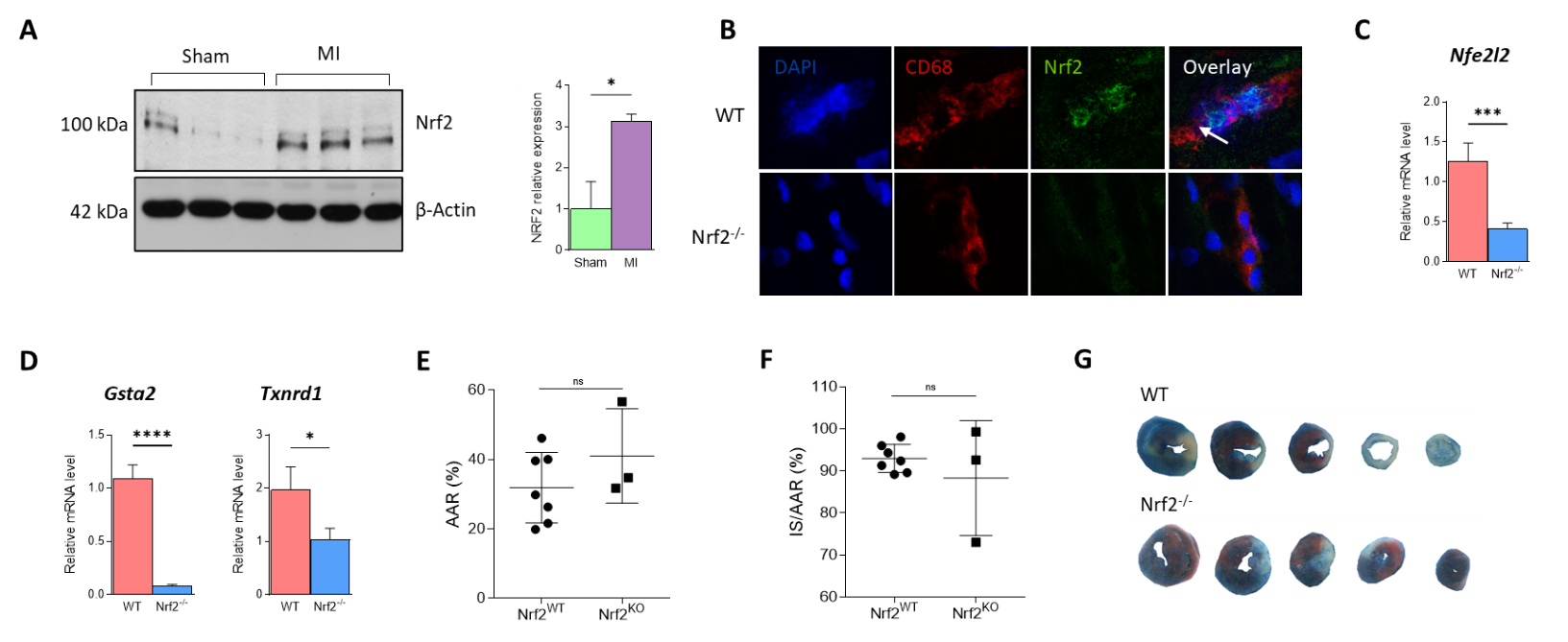

#### Supplemental Fig 2: No significant differences observed in blood leucocytes of global Nrf2^-/-^ mice after MI

9-13 week old mice were anaesthetised and subjected to permanent LAD ligation prior to recovery for 72 h**.** Figures show quantification of total leukocytes (defined as CD45^+^) and subsets, defined as CD45^+^CD19^-^Ly6G^-^Ly6C^hi^ (M1) and CD45^+^CD19^-^Ly6G^-^Ly6C^lo^ (M2) monocytes, CD45^+^CD19^-^Ly6G^+^CD206^-^ (N1) and CD45^+^CD19^-^Ly6G^-^CD206^-^ (N2) neutrophils. Data shown as cells/ml blood, n=7-11 per group compared using 1-way ANOVA with Tukey’s post-test, all non-significant.

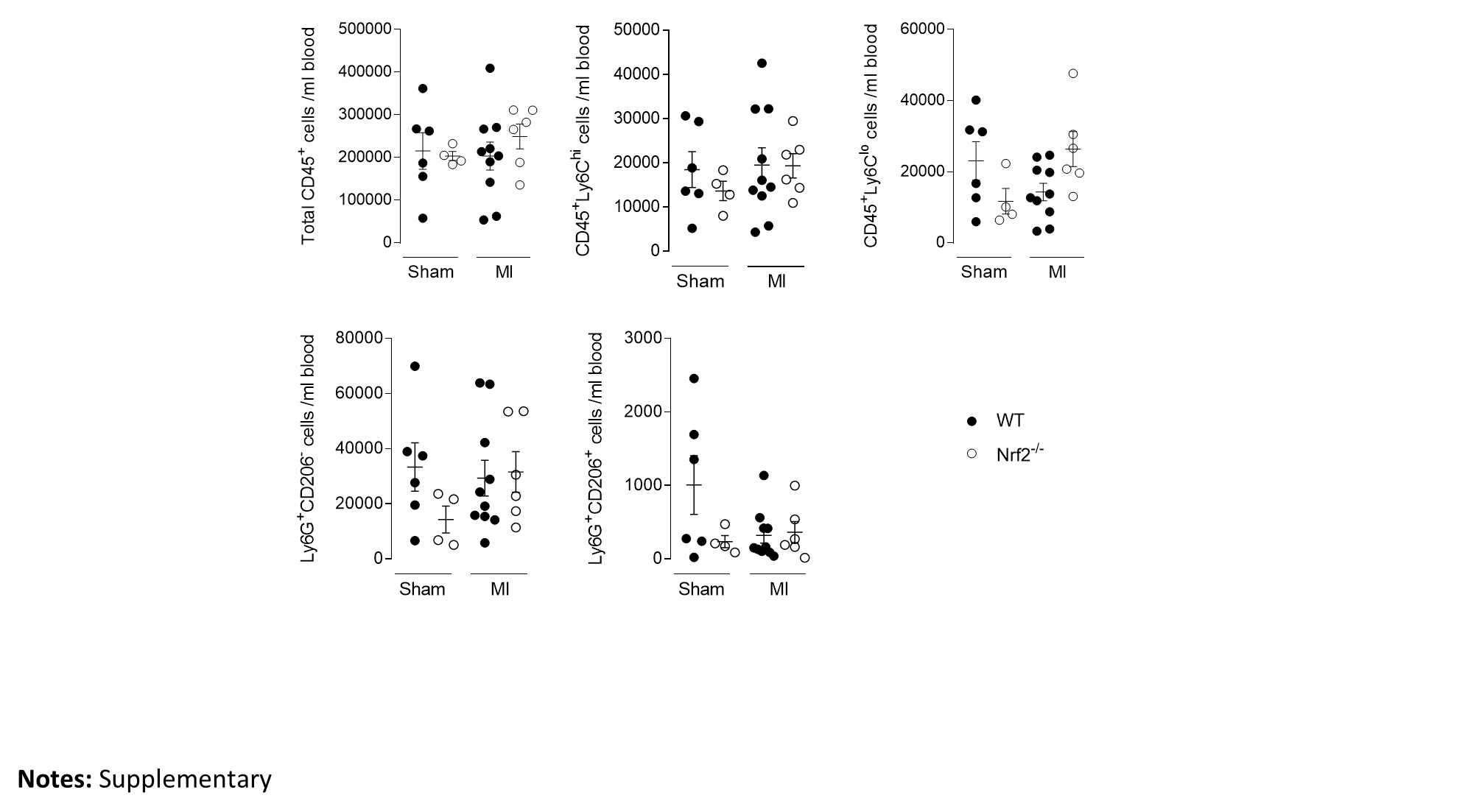

#### Supplemental Fig 3: Integrated clusters

Integrated heatmap showing clusters on x axis and marker genes on y axis. Yellow and purple indicate high and low expression, respectively.

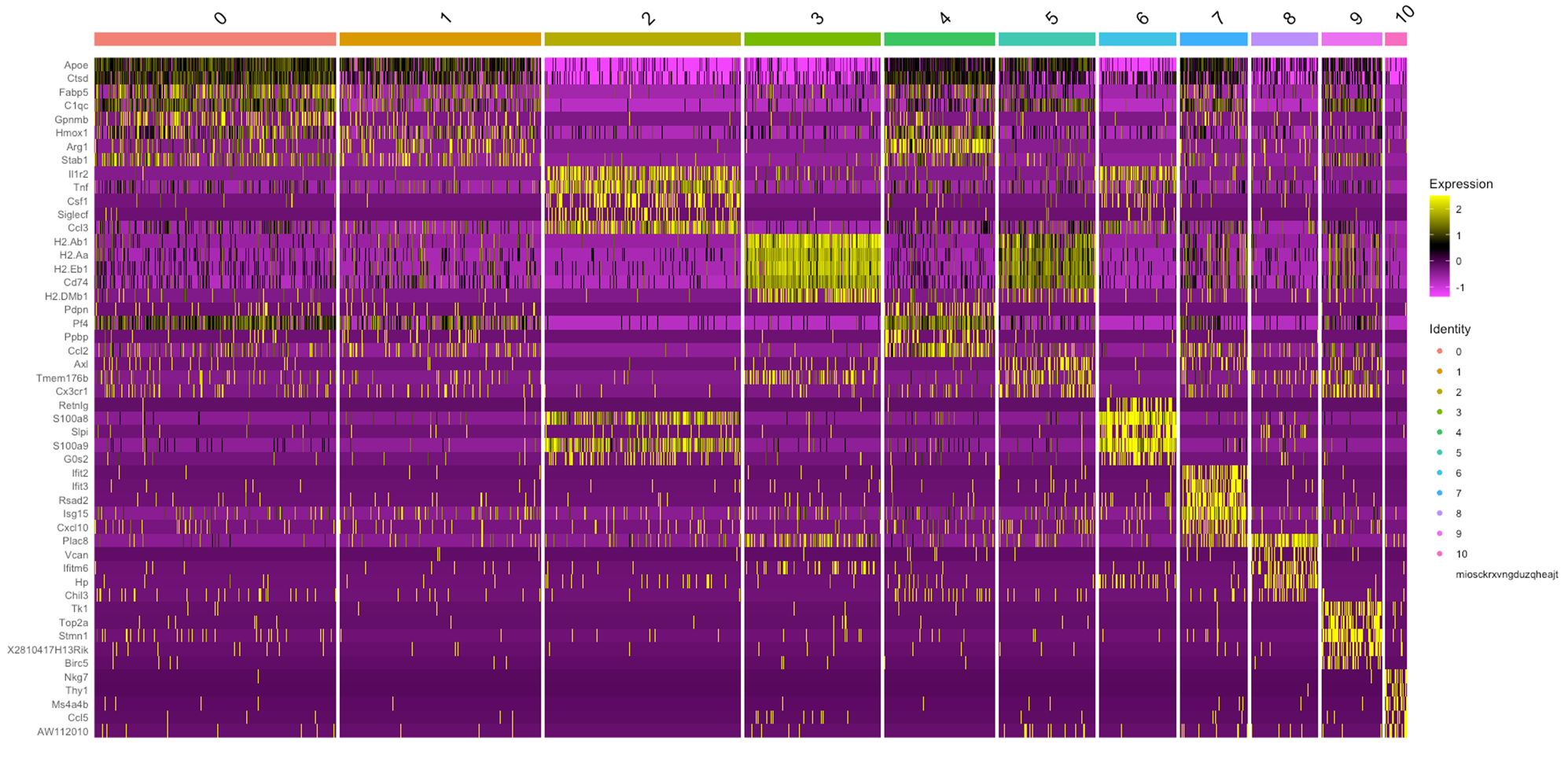

#### Supplemental Fig 4: ImmGen analysis of cluster 0 (macrophage 1)

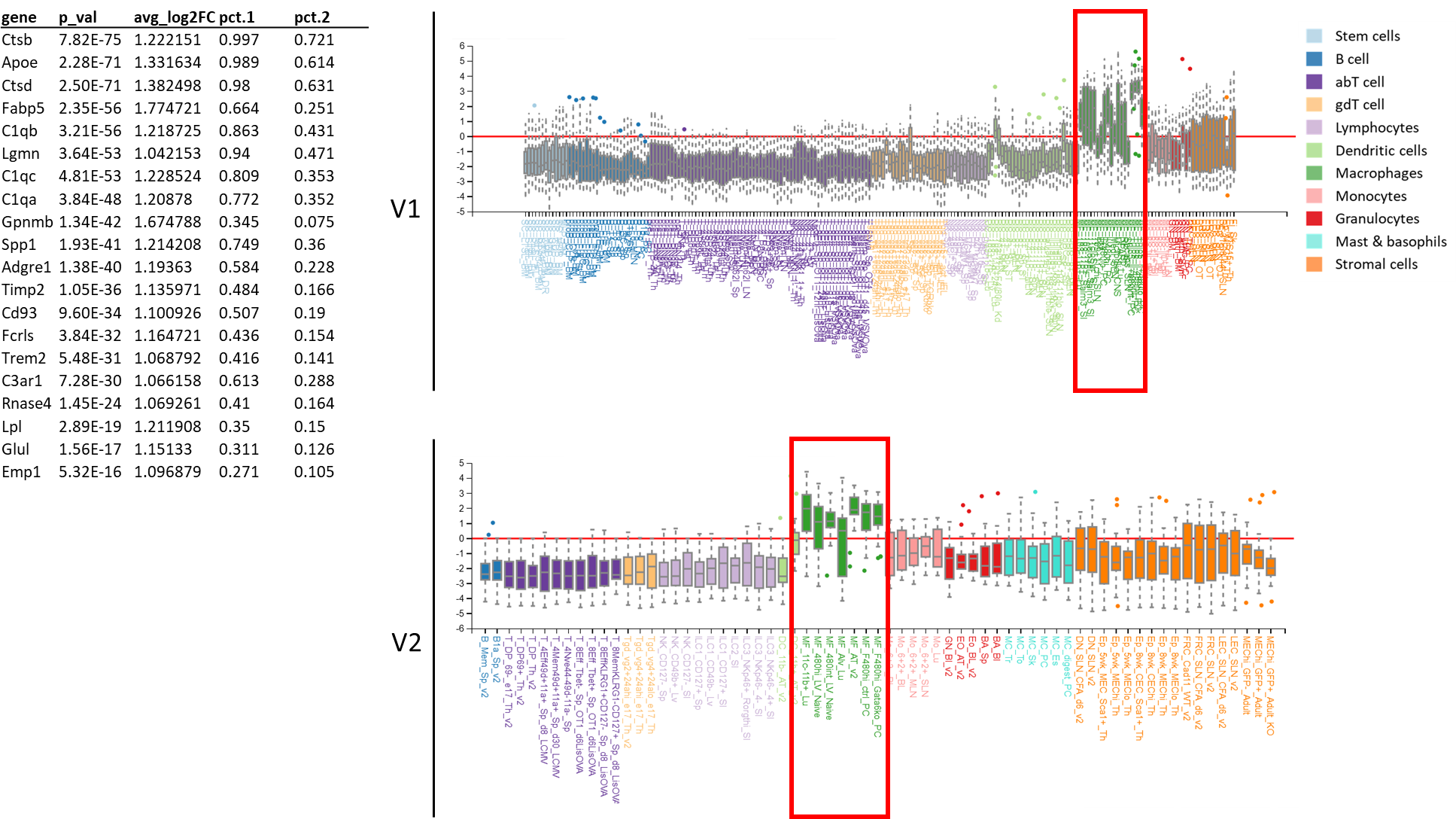

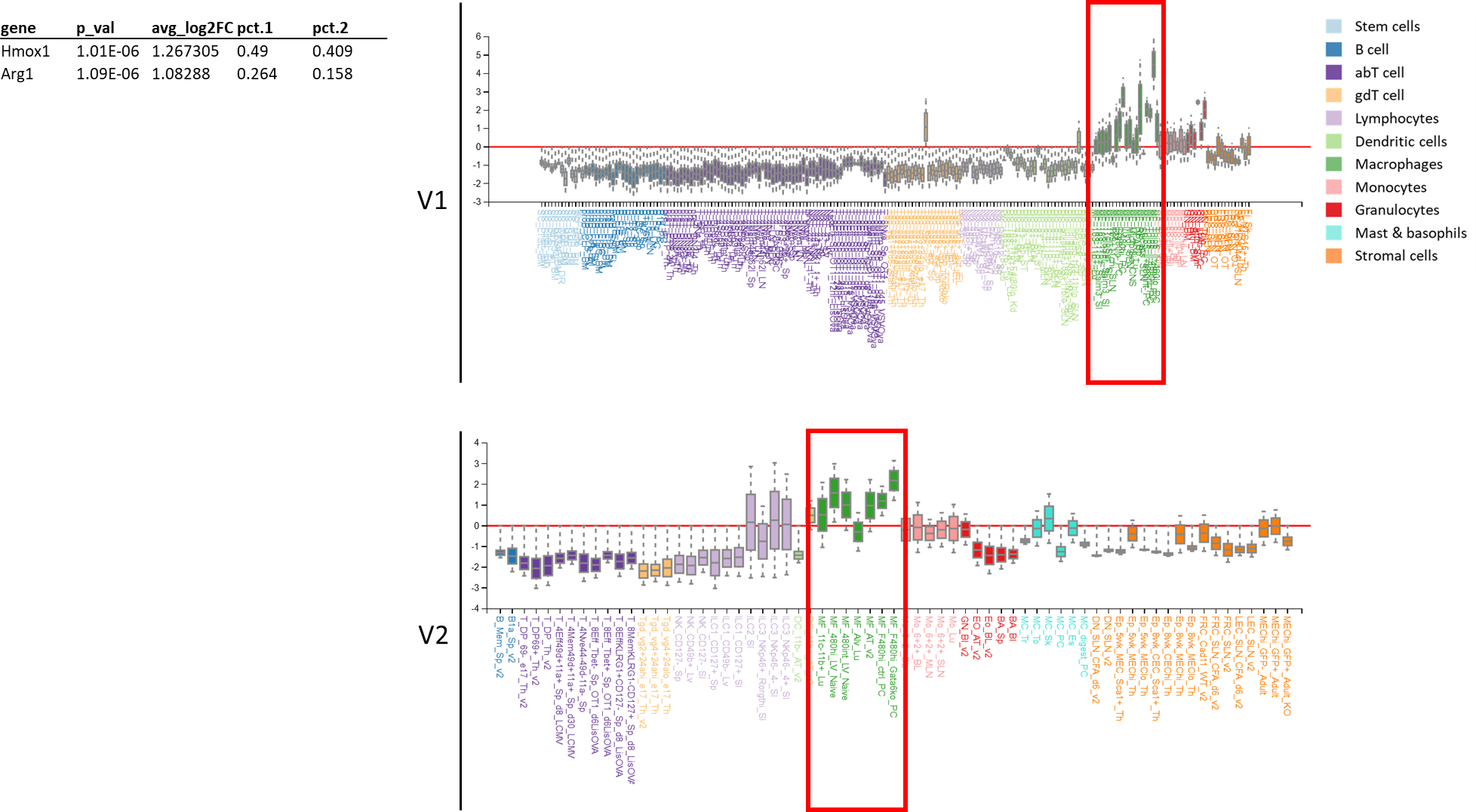

#### Supplemental Fig 5: ImmGen analysis of cluster 1 (macrophage 2)

#### Supplemental Fig 6: ImmGen analysis of cluster 2 (granulocyte 1)

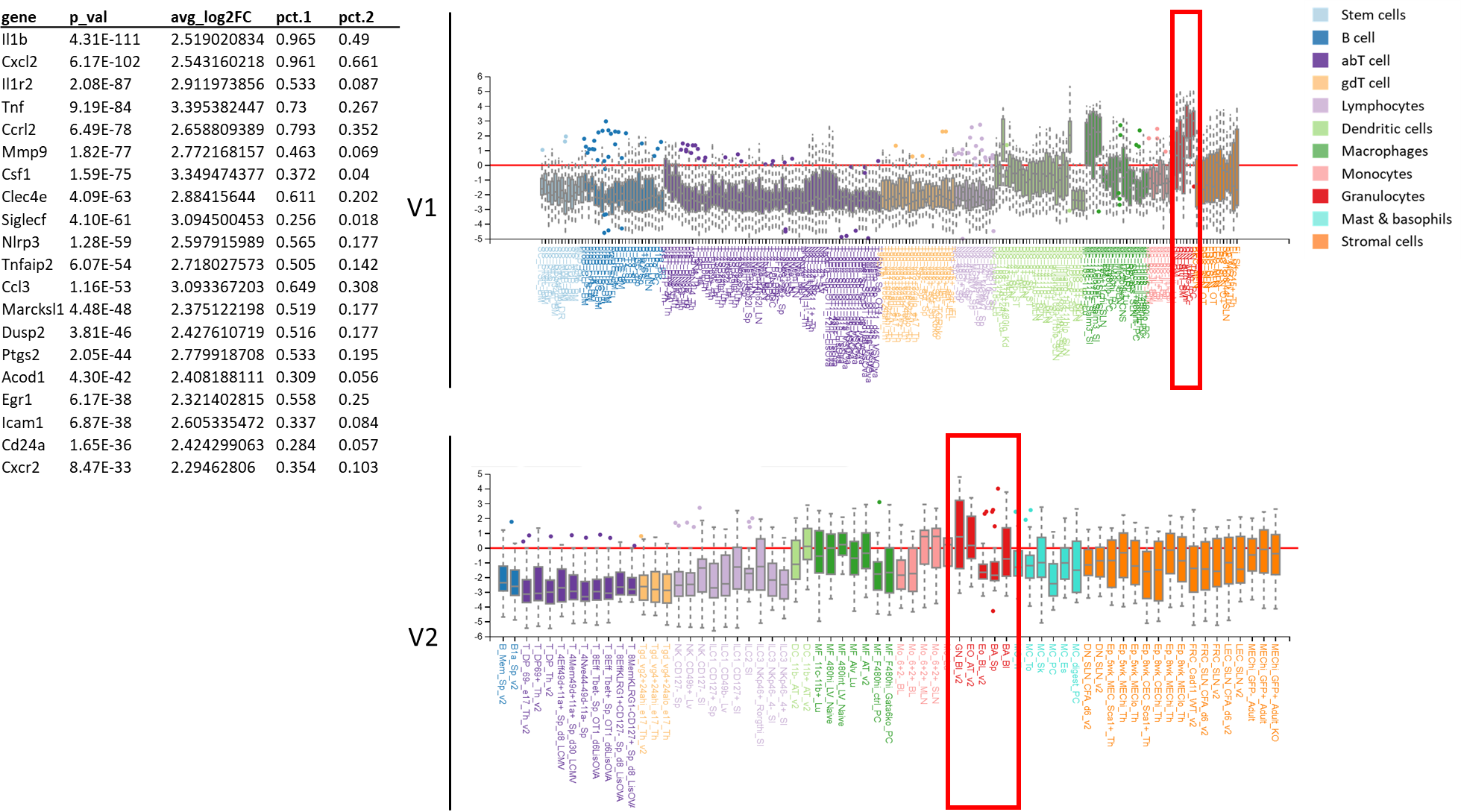

#### Supplemental Fig 7: ImmGen analysis of cluster 3 (macrophage 3)

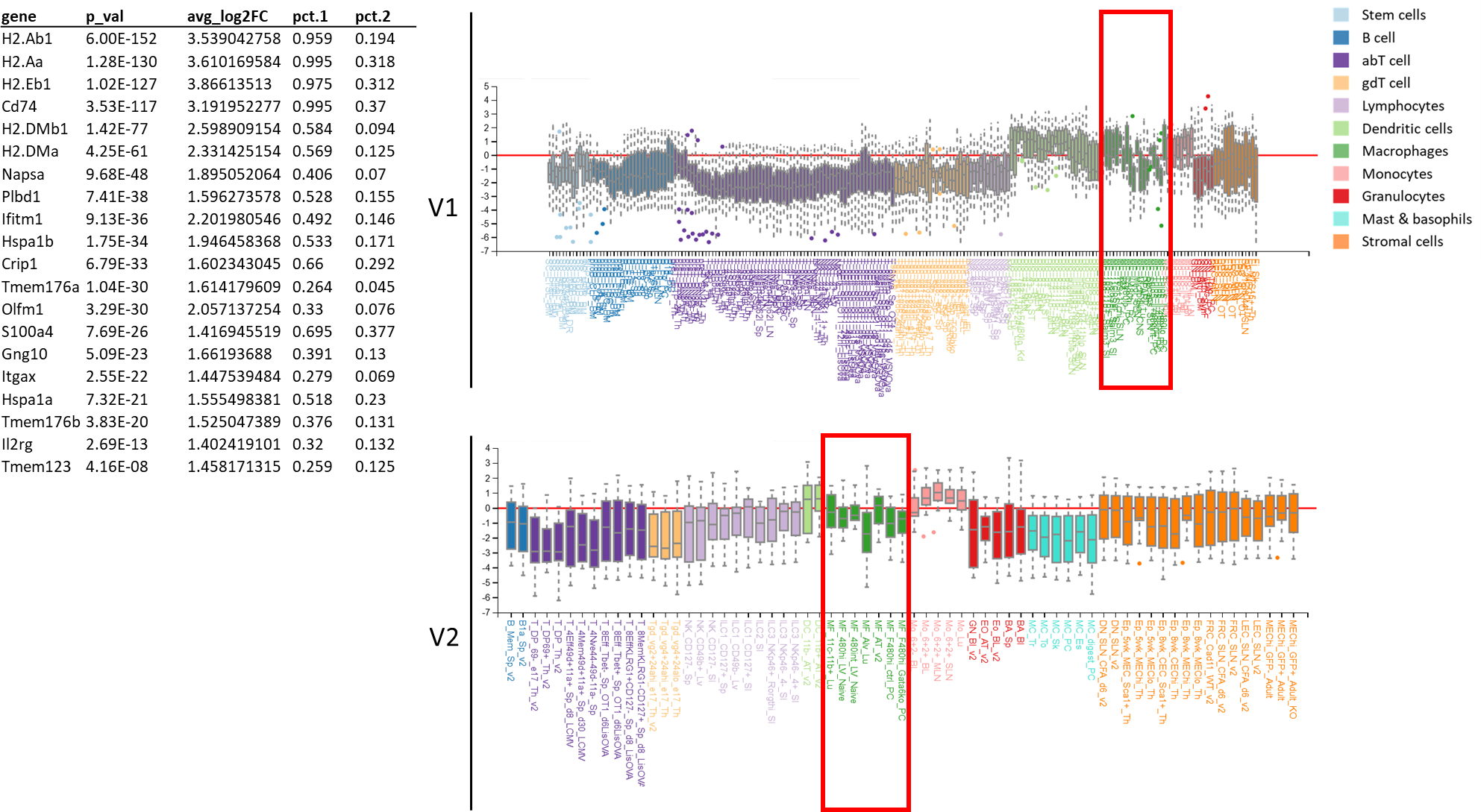

#### Supplemental Fig 8: ImmGen analysis of cluster 4 (macrophage 4)

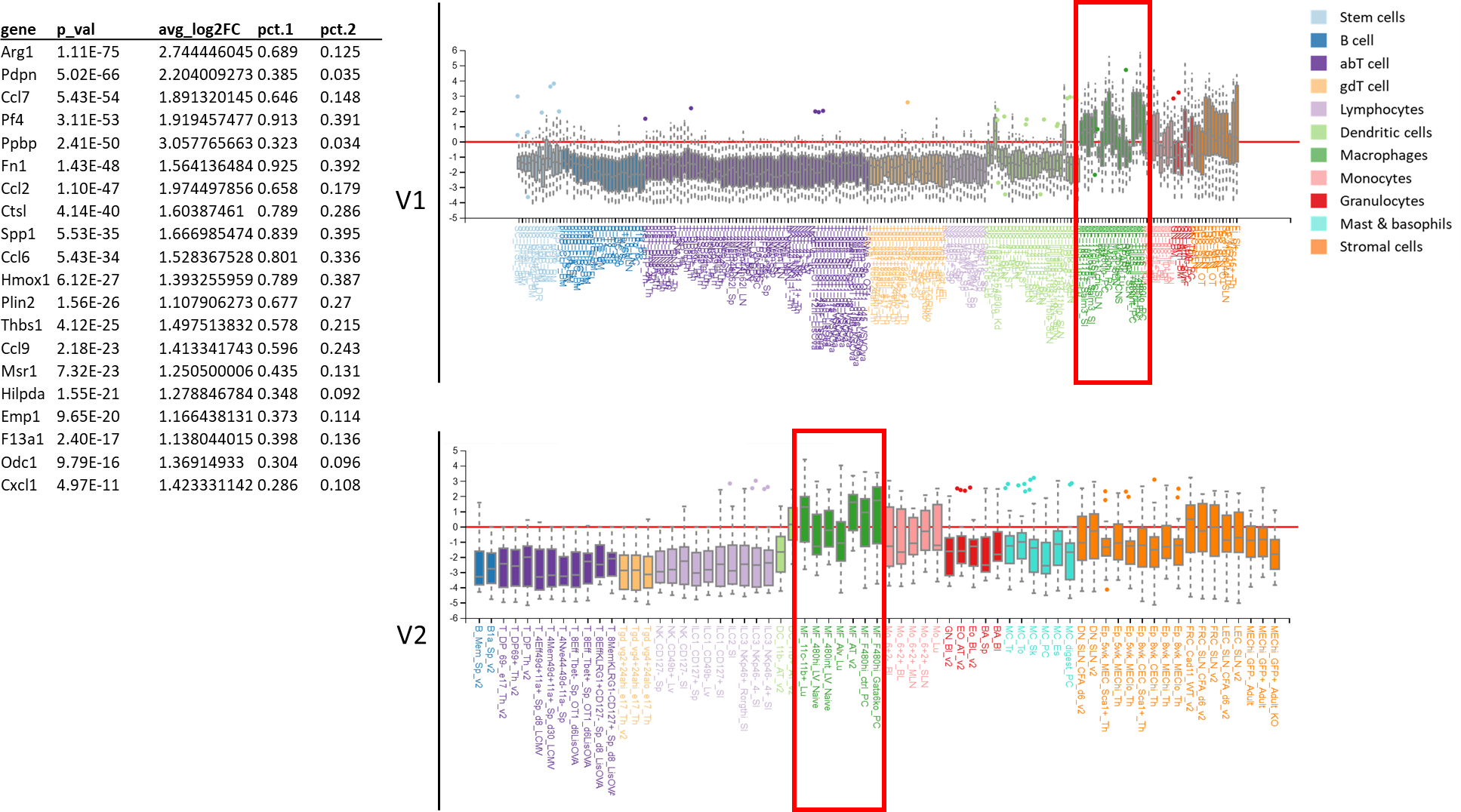

#### Supplemental Fig 9: ImmGen analysis of cluster 5 (dendritic cell)

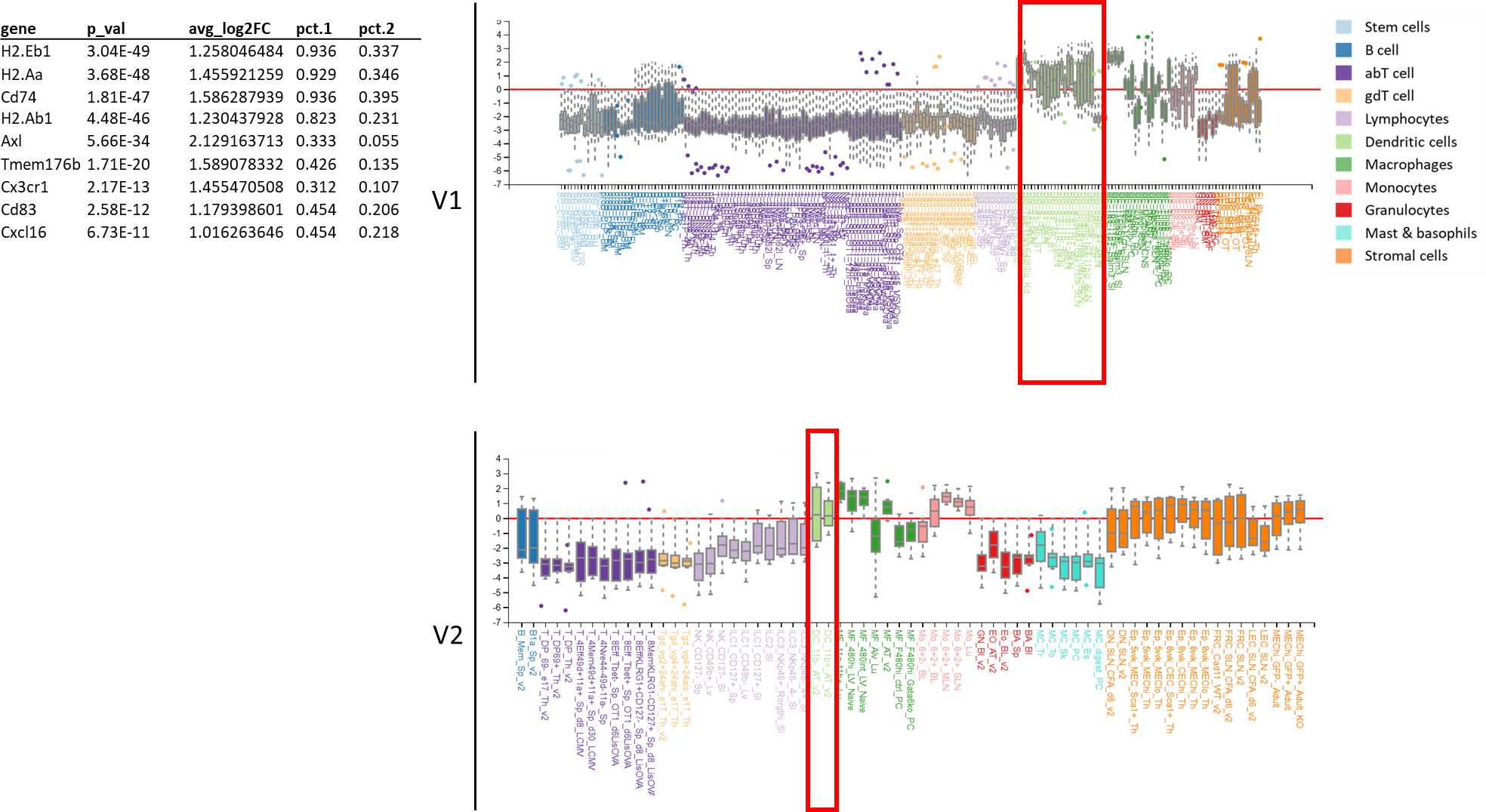

#### Supplemental Fig 10: ImmGen analysis of cluster 6 (granulocyte 2)

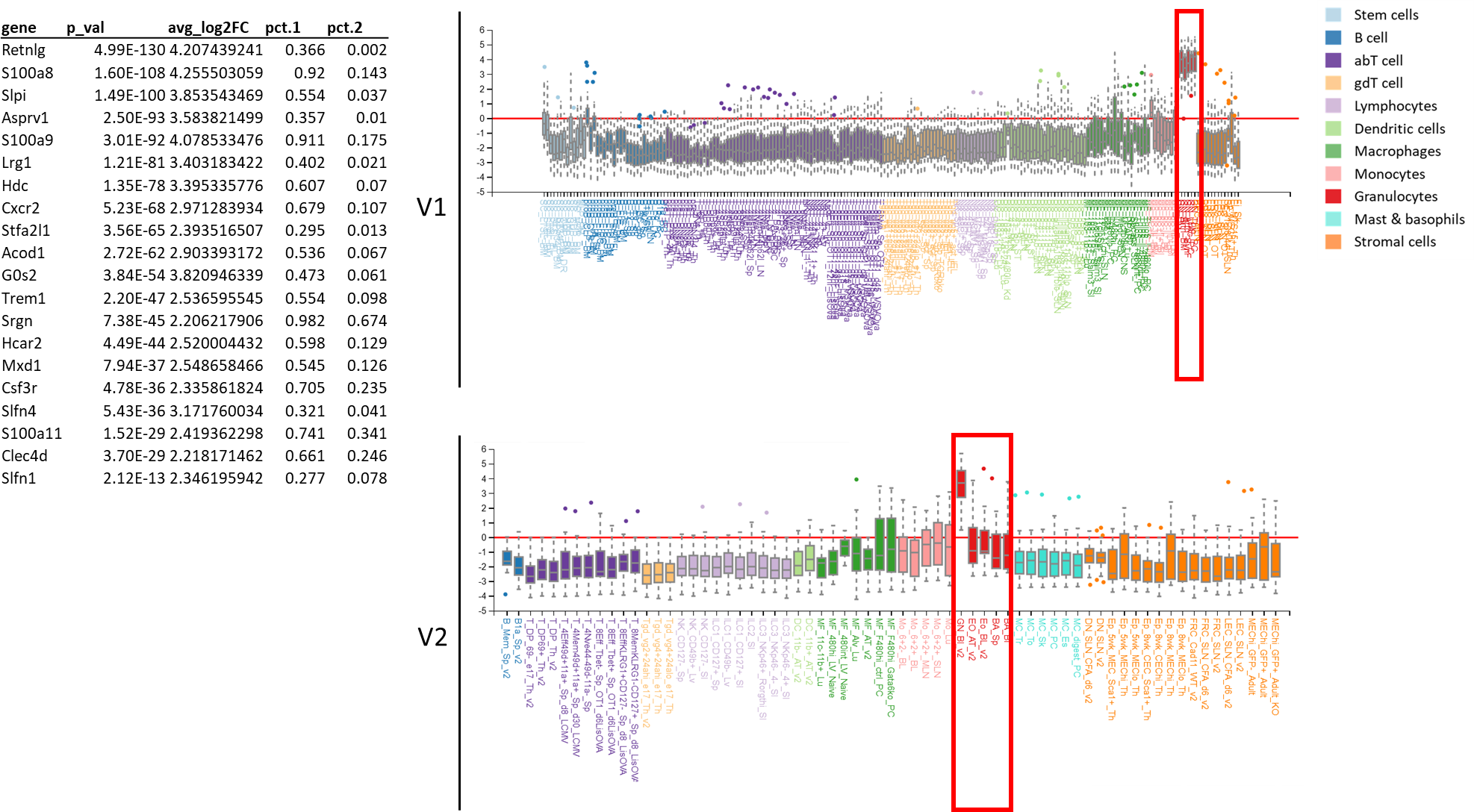

#### Supplemental Fig 11: ImmGen analysis of cluster 7 (macrophage 5)

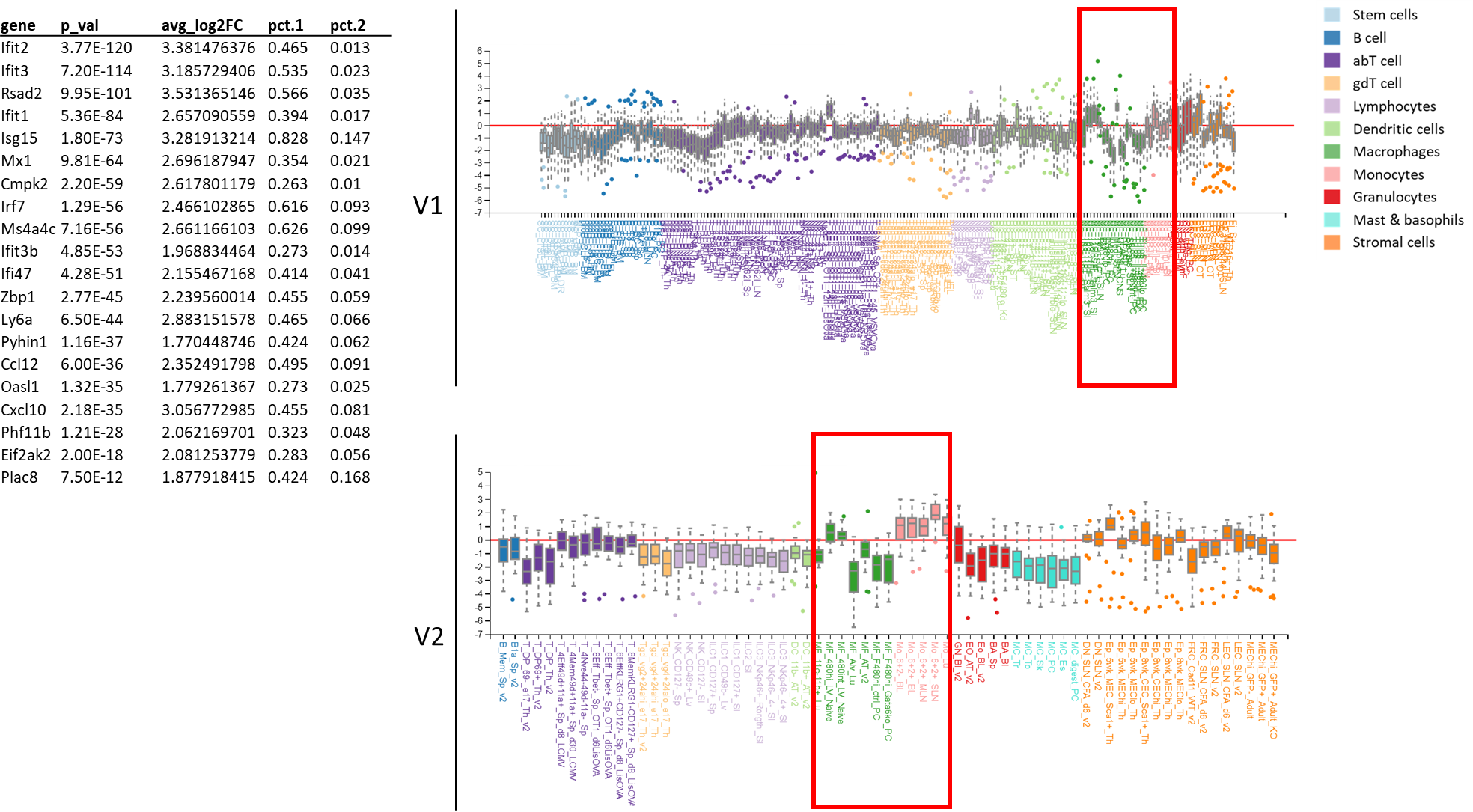

#### Supplemental Fig 12: ImmGen analysis of cluster 8 (monocyte)

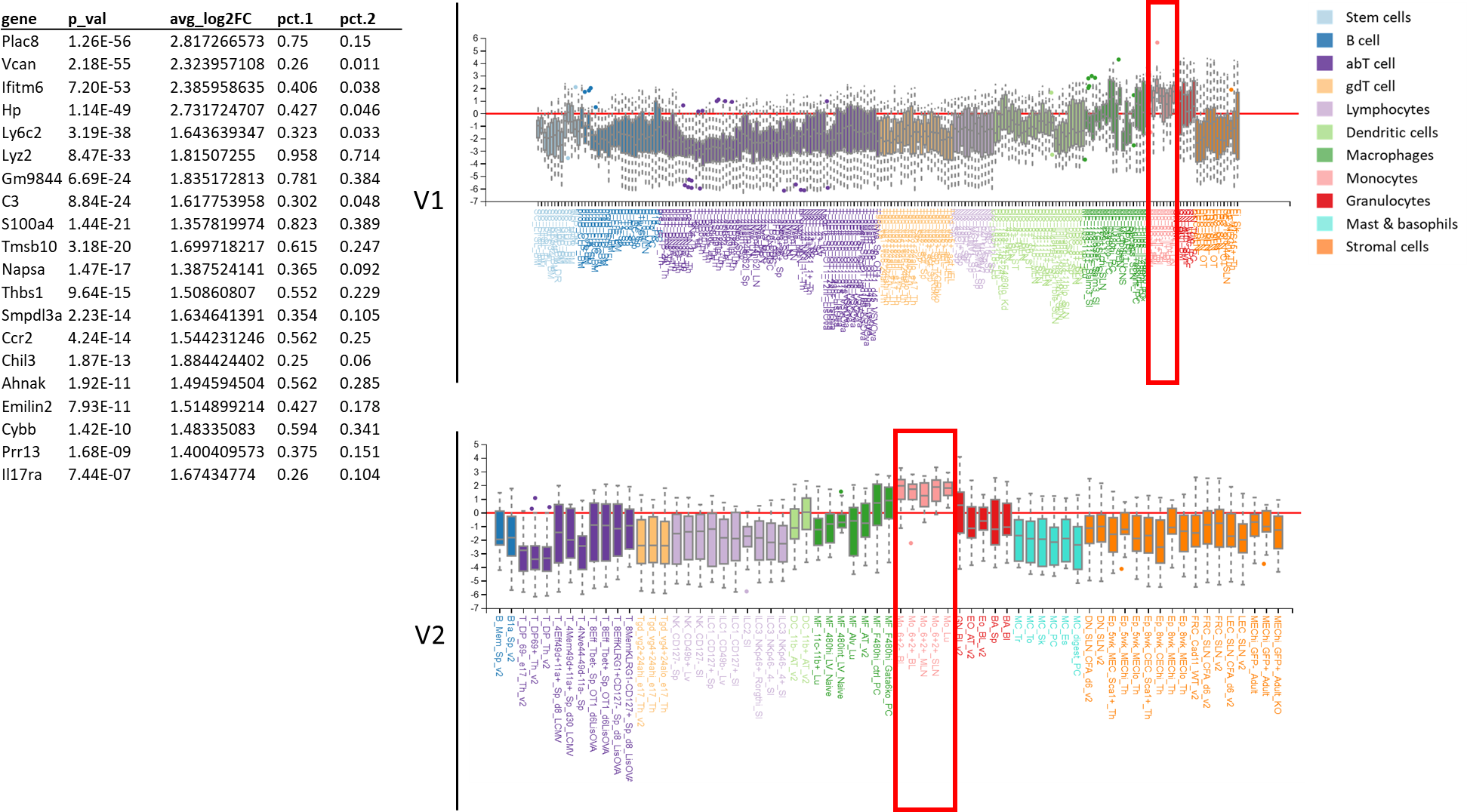

#### Supplemental Fig 13: ImmGen analysis of cluster 9 (proliferating)

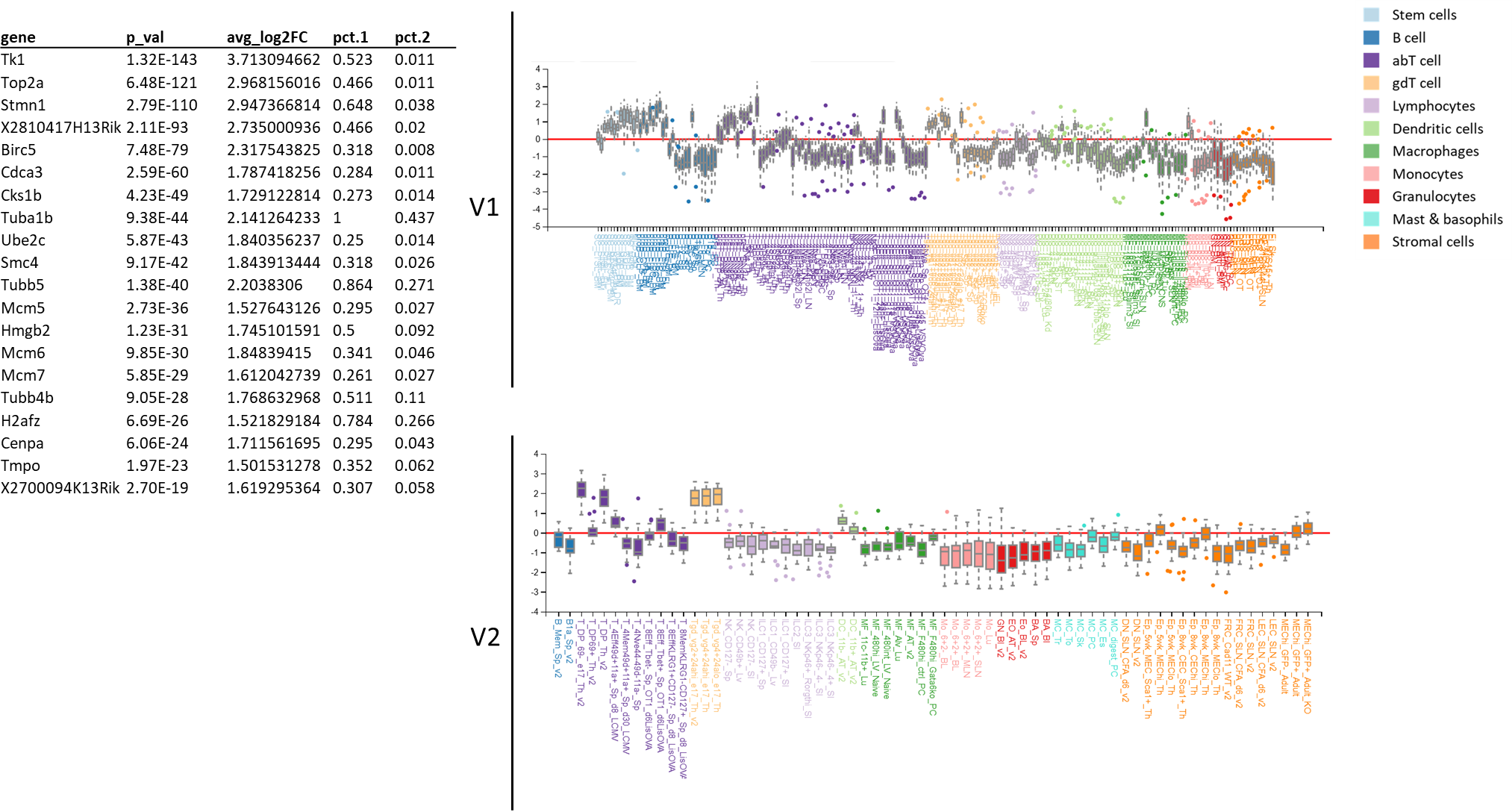

#### Supplemental Fig 14: ImmGen analysis of cluster 10 (T / NK cell)

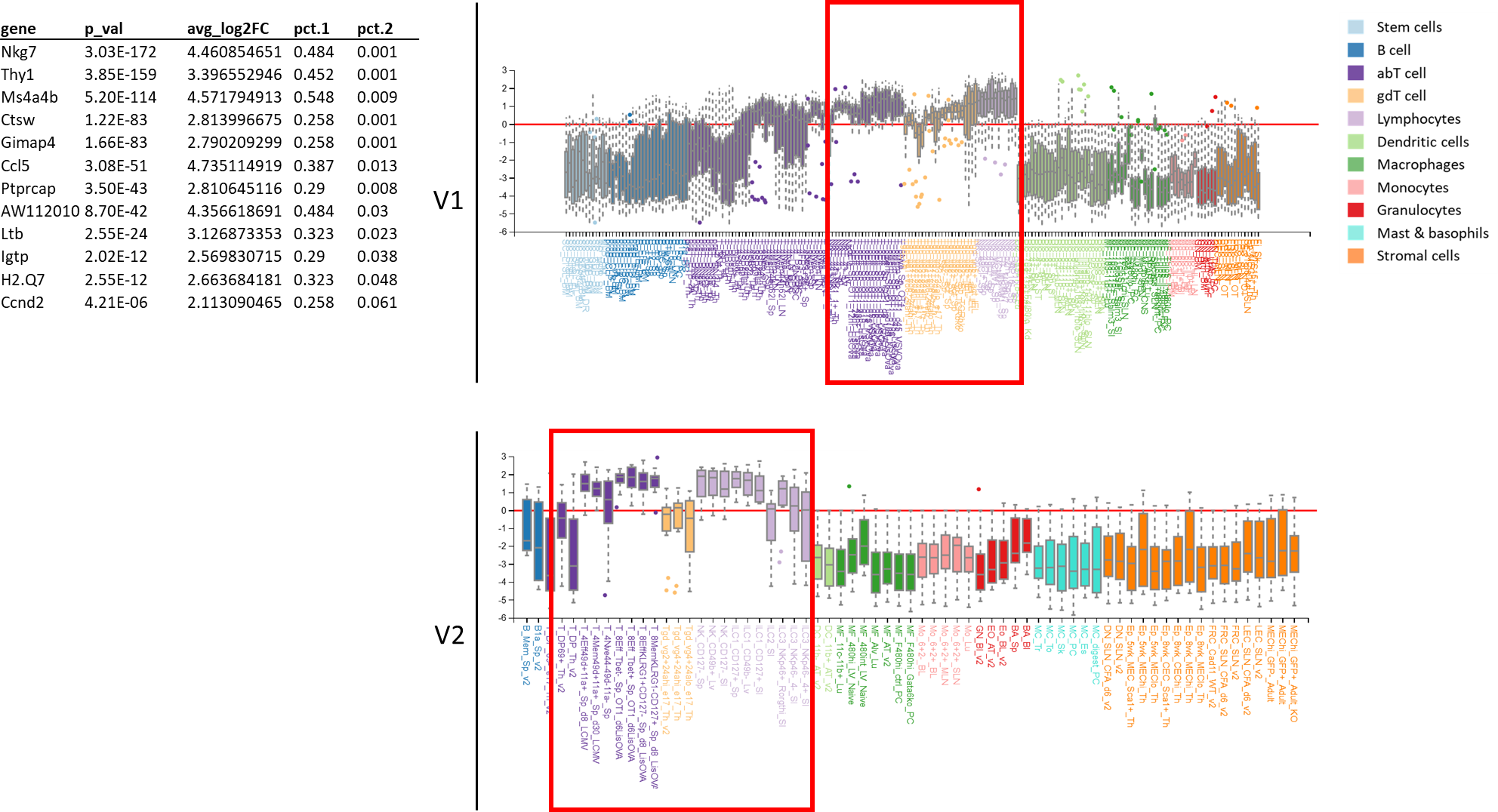

#### Supplemental Fig 15: Nrf2-stimulated genes (NRGs) after MI

Clustered, publicly available single-cell RNA-seq data for 1,858 single cells isolated from a single WT mouse 4d after permanent LCA occlusion. Violin plots show the expression probability distributions of NRGs in each cell cluster.

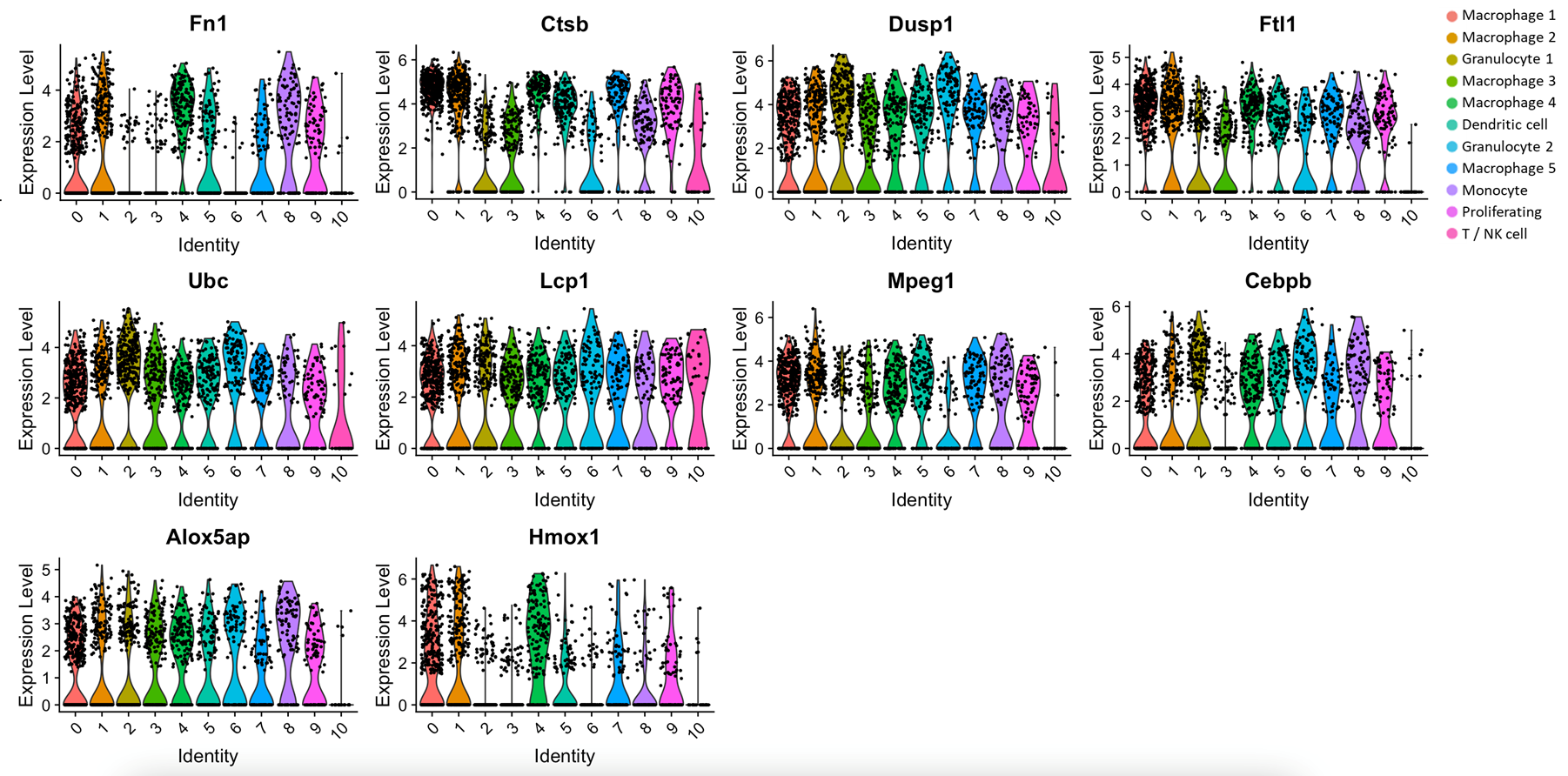

### Supplemental tables

#### Supplemental table 1: Primer sequences

| **Target** | **Forward (3’->5’)** | **Reverse (3’->5’)** |
| --- | --- | --- |
| Actb | CGTGAAAAGATGACCCAGATCA | TGGTACGACCAGAGGCATACAG |
| Gapdh | CCTAGACAAAATGGTGAAG | GACTCCACGACATACTCAGC |
| Canx | TGCCAGAAACAGCTCCTCAT | AGTTTGGTGACGATGCCTTT |
| Ccl2 | TGATCCCAATGAGTAGGCTGGAG | ATGTCTGGACCCATTCCTTCTTG |
| Ccr2 | ACCTGTAAATGCCATGCAAGT | TGTCTTCCATTTCCTTTGATTTG |
| Cxcl2 | AGACAGAAGTCATAGCCACTCTCAAG | CCTCCTTTCCAGGTCAGTTAGC |
| Cxcr4 | TCAGTGGCTGACCTCCTCTT | CTTGGCCTTTGACTGTTGGT |
| Cxcr2 | AGCAAACACCTCTACTACCCTCTA | GGGCTGCATCAATTCAAATACCA |
| Hmox1 | CAGCCCCACCAAGTTCAAA | TCAGGTGTCATCTCCAGAGTG |
| Il10 | GGGTCTTGGGAAGAGAAACC | CATTCCCAGAGGAATTGCAT |
| Il6 | GAGGATACCACTCCCAACAGACC | AAGTGCATCATCGTTGTTCATACA |
| Sod2 | CACACATTAACGCGCAGATCA | GGTGGCGTTGAGATTGTTCA |
| Prdx3 | CCAGAGTCCCCTACGATCAA | TCAAGGCATTGGAAGGATTG |
| Nfe2l2 | CTACTCCCAGGTTGCCCACA | CGACTCATGGTCATCTACAAATGG |
| Gsta2 | GCTTGATGCCAGCCTTCTG | GGCTGCTGATTCTGCTGCTCTTGA |
| Txnrd1 | GATGCACCAGGCAGCTTTG | TCTTCGACTTTCCAGCCATAGT |
| Col4a1 | AAAGGCTCTCCGGGTTCAAT | GCCGATGTCTCCACGACTAC |
| Dcn | AACTGTGCTATGGAGTAGAAGCA | ATCTCATGTATTTTCACGACCTTTT |
| Lamc1 | GCAGCCCTGTTGGTTCTCTC | TTACACTCGTCTGTGCTGCC |
| Gstm2 | GAAAGCACAACCTGTGTGGAGAG | AGCTTCATCTTCTCAGGGAGAC |

#### Supplemental table 2: Antibodies used for FACS experiments

| **Marker** | **Fluorochrome** | **Concentration** | **Supplier** | **Catalogue number** |
| --- | --- | --- | --- | --- |
| CD45 | APC Cy7 | 1:200 | BioLegend | 103115 |
| CD19 | BV395 | 1:100 | BD Biosciences | 536557 |
| CCR2 | PE | 1:100 | BioLegend | 150609 |
| Ly6G | BV785 | 1:200 | BioLegend | 127645 |
| Ly6C | BV605 | 1:100 | BioLegend | 128035 |
| CD11b | BV510 | 1:200 | BioLegend | 101245 |
| MHC II | BV711 | 1:1000 | BioLegend | 107643 |
| IL6 | APC | 1:100 | BioLegend | 504507 |
| iNOS | FITC | 1:100 | BD Biosciences | 610330 |
| CD206 | BV421 | 1:100 | BioLegend | 141717 |
| F4/80 | PE Cy7 | 1:100 | eBioscience | 25-4801-82 |

Supplemental referencesReference List
